# iSeqQC: A Tool for Expression-Based Quality Control in RNA Sequencing

**DOI:** 10.1101/768101

**Authors:** Gaurav Kumar, Adam Ertel, George Feldman, Joan Kupper, Paolo Fortina

**Author notes:** **Correspondence:** Gaurav Kumar, PhD, Department of Cancer Biology, BLSB 1009 233 South 10th Street Philadelphia, PA-19107, USA.

## Abstract

Quality Control in any high-throughput sequencing technology is a critical step, which if overlooked can compromise the data. A number of methods exist to identify biases during sequencing or alignment, yet not many tools exist to interpret biases due to outliers or batch effects. Hence, we developed iSeqQC, an expression-based QC tool that detects outliers either produced by batch effects due to laboratory conditions or due to dissimilarity within a phenotypic group. iSeqQC implements various statistical approaches including unsupervised clustering, agglomerative hierarchical clustering and correlation coefficients to provide insight into outliers. It can be utilized either through command-line (Github: https://github.com/gkumar09/iSeqQC) or web-interface (http://cancerwebpa.jefferson.edu/iSeqQC). iSeqQC is a fast, light-weight, expression-based QC tool that detects outliers by implementing various statistical approaches.

## INTRODUCTION

High-throughput experiments are complex and prone to numerous biases during sample preparation, library preparation and sequencing. Therefore, Quality Control (QC) is critical and if overlooked, can compromise the data. To reduce false discoveries from any quantitative sequencing experiment such as RNA-seq, miRNA-seq and ATAC-seq, QC can be categorized in three different phases. In phase one, quality of raw read sequences is analyzed to detect bad quality bases. This is mainly performed on raw FASTQ files using tools including FastQC [1], FASTX-Toolkit [2], NGS QC Toolkit [3], and PrinSeq [4]. In phase two, mapping quality, read count distribution, mean insert size distribution, mean depth distribution, base quality and capture efficiency are observed on aligned BAM files to detect sample biases occurring during library preparation. This is mainly done using tools like RseQC [5], RNA-SeQC [6], QC3 [7] and QoRTs [8]. In phase three, sample heterogeneity, outliers and any cross-sample contamination are detected using various statistical approaches such as correlations and dimensional reductions. There is neither any defined rule to study QC in the third phase nor is any specific QC tool available to study in-depth sample quality.

Here, we present iSeqQC- an expression-based quality control tool to detect outliers either produced by batch effects due to laboratory conditions, reagent lots, personnel differences, different experiment times, or merely due to dissimilarity within a phenotypic group. Very straight-forward to use, iSeqQC uses count matrix data from an expression analysis to produce QC metrics in the form of graphical plots defining relationships of all samples.

## METHODS

### iSeqQC algorithm

iSeqQC provides comprehensive information to identify any outliers in the sequencing experiment due to any technical biases. Developed in R, it can be utilized through web-interface (http://cancerwebpa.jefferson.edu/iSeqQC), running the Shiny Server program, and also through command-line (GitHub, https://github.com/gkumar09/iSeqQC).

iSeqQC requires two tab-delimited text files to execute: 1) a sample phenotype file with information on sample names and phenotypes; 2) an unnormalized count matrix from any read summarization tool such as RSEM [9], HTseq [10], feature counts [11] and so on. Using information from sample phenotype file, iSeqQC first matches the sample names to the count matrix, then implements the following statistical approaches to provide comprehensive QC metrics:

1. Summary statistics and counts distribution: it uses unnormalized expression matrix to provide basic descriptive summary statistics and normally distributed unnormalized expression matrix to provide counts distribution per sample.
2. Mapped reads density: after Reads Per Million (RPM) normalization using the following formula, density of mapped reads is estimated for each sample

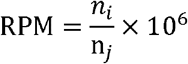

where n_i_ is number of reads mapped to a gene and n_j_ is library size
3. Housekeeping gene expression: Expression profile of two housekeeping genes, GAPDH and ACTB for all samples.
4. Principal Component analysis (PCA): After z-transforming the expression matrix so that each row has a mean of 0 and a variance of 1, PCA, a dimensionality reduction algorithm is implemented to linear transform the data and observe the variance between samples. Z-score normalization is performed using

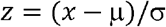

where μ is mean and σ is variance
5. Hierarchical relationship: Measuring the distance of similarity between samples, hierarchical clustering is also implemented on the z-transformed expression matrix.
6. Correlation: to detect strength of association between the samples, Pearson (quantity-based) and Spearman (rank-based) correlation are implemented on the normally distributed unnormalized expression matrix.

After successfully executed, iSeqQC provides QC metrics in the form of a table and several graphical plots. First, it uses raw expression data to provide descriptive statistics, output in a form of a ‘summary statistics’ table. For each sample, it provides number of detected genes, mean expression, standard deviation, median expression, minimum expression, maximum expression, range of expression, skewness (symmetry of expression distribution) and kurtosis (tails of distribution). Next, the expression data is z-score normalized to obtain normal distribution and then displayed as a ‘count-distribution’ box-plot to provide overall distribution of the expression of each sample with minimum, maximum and median expression. Further, the raw expression data is RPM normalized and density distribution of mapped reads is provided in a form of ‘mapped read density’ plot to observe any sample with no or low expressing reads. Due to stable expression of housekeeping genes, they are often used in sequencing experiments to normalize mRNA levels between different samples. iSeqQC uses GAPDH and ACTB expression data to detect whether samples show high expression of these two genes. All this information is used as a first sign to detect any outlier and could be further investigated for any possible biasness.

After implementing basic statistics algorithms, iSeqQC implements various dimensionality reduction approaches to extract any technical bias in the sequencing experiment. Here, it first z-score normalizes the expression data and implements PCA unsupervised clustering method to identify the principal directions or variations called as components. The first two principal components which mainly are the highest source of variance are then displayed as a plot. This plot further segregates the samples based on their phenotype (data obtained from sample phenotype sheet). Next hierarchical clustering, another method of dimensionality reduction is implemented to provide the distance of similarity between replicates in a specific phenotypic group. Using the Agglomerative method, it assigns each sample to its own cluster and then computes distance between each cluster and joins the two most similar clusters together. Next, iSeqQC utilizes correlation coefficients to detect the strength and direction of the relationship between the samples. It uses Pearson correlation, which evaluates the linear relationship between the samples and Spearman correlation, which is a rank-based method that can range from −1 to +1. The direction of the relationship is indicated by the value of the coefficient; samples with a close relationship tend to be in the positive range and vice versa. Output in the form of a plot, all this comprehensive information cumulatively provides sufficient indication of any outlier sample or cross-sample contamination.

### Sample Collection

Surgical samples from Depuytrens affected patients and controls were collected and stored in RNAlater (ThermoFischer Scientific, MA, USA, catalogue no.-AM7020) at −80°C. Under sterile conditions, total RNA was extracted using the Qiagen miRNeasy mini-kit (Qiagen, MD, USA, catalogue no.-217004).

### RNA Extraction, Library Preparation and Sequencing

4 ng of total RNA was used to prepare libraries using the Takara Bio SMARTer Stranded Total RNA-Seq Kit (Takara Bio, CA, USA, catalogue no.-634837) following manufacturer’s protocol. The final libraries were sequenced on NextSeq 500 using 75 bp paired-end chemistry.

### Sequencing and library QC

Sequencing QC to obtain any read errors, poor quality reads and primer or adapter contamination was observed using FastQC. Inconsistencies in sample and library preparation was observed using QC3 [7], QoRTs [8], and RSeqQC [5].

### Differential expression analysis

Differential gene expression was performed using diseased and control samples using the DESeq2 [12] package in R/Bioconductor. Genes were considered differentially expressed (DE) if they had adjusted p value ≤ 0.05 and absolute fold change ≥ 2. All plots were constructed using R/Bioconductor.

## RESULTS

To demonstrate the importance of expression QC and performance of iSeqQC, we utilized RNA-seq samples sequenced in our laboratory to study Depuytrens disease.

Following our laboratory standard protocols, all samples were tested first to access integrity of RNA (RIN) score and were within range of the requirements of the library preparation kit (≥ 2). The disease samples had a RIN score between 4-5 and control samples were between 2-3.

Next, samples were sequenced and resultant FASTQ files were examined for sequencing errors using FastQC in phase one of QC. Here, for all samples, per base sequence quality for all bases at each position was observed to be >30, demonstrating a base call accuracy >99.9% **(Fig. 1a)**. Additional metrics such as per base sequence and GC content also were observed to be good quality.

**Fig 1.**
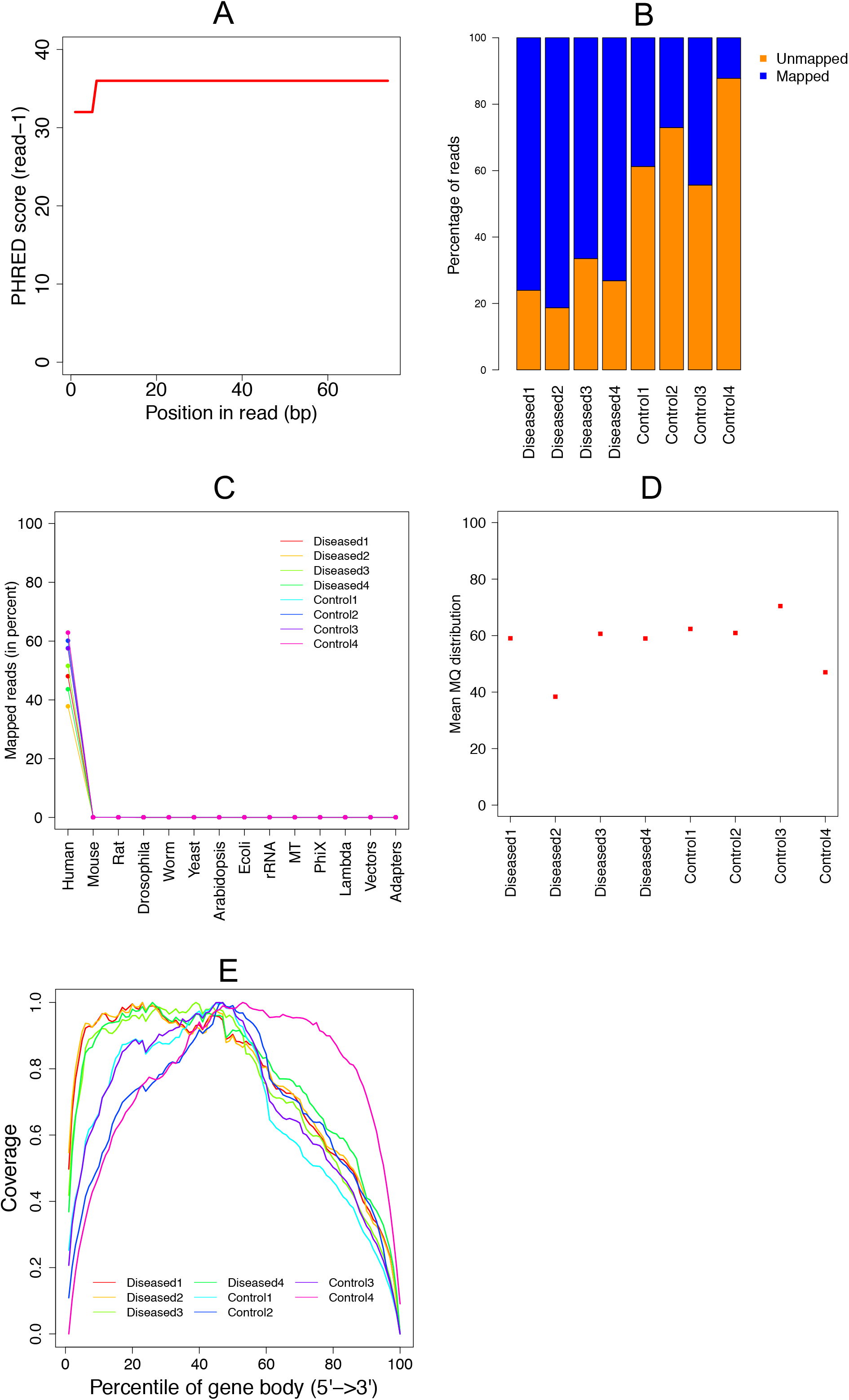
Quality control metrics using existing tools. A) Per base sequencing quality averaged for all diseased and control samples; B) Mapping statistics showing percentage of mapped versus unmapped reads in diseased and control samples; C) Percentage of reads mapped uniquely to human genome demonstrating no contamination in the libraries; D) Average mapping quality showing no outlier; E) Coverage uniformity over gene body for all samples showing no outlier.

In phase two, QC was observed on the aligned BAM files using QC3 and RSeqQC. QC3 was used to observe mapping statistics, where all diseased samples had >70% reads mapped to the reference genome showing high quality RNA samples. However, control samples had an overall low mapping percentage (control1, 2, 3: ~40% and control4: ~20%) as shown in **Fig. 1b**. With the suspicion of DNA or any other contamination, we used FastQ Screen to further investigate the low mapping of control samples. A high proportion of mapped reads were mapping to only the human genome **(Fig. 1c)**. Next, RSeqQC was used to observe average mapping quality, where diseased2 and control4 were observed to have a low but acceptable quality as shown in **Fig. 1d**. Additionally, the coverage over gene body analysis also showed an acceptable coverage uniformity over gene body in all samples **(Fig. 1e)**. These existing sequencing and library QC tools were inconclusive to detect any outliers in the study.

Next, in phase three, QC was observed on expression data using iSeqQC, which generates a summary table and 7 different plots to infer QC. Upon investigating the Principal Component Analysis (PCA) clustering **(Fig. 2a)**, hierarchical clustering **(Fig. 2b)**, Pearson **(Fig. 2c)** and Spearman clustering **(Fig. 2d)** from iSeqQC output, we observed tight clustering and correlation of each sample in its phenotypic group, hence no biases. However, the ‘housekeeping gene’ plot showed an overall low expression of ACTB and GAPDH in control4 sample when compared to other samples tested **(Fig. 2e)**. Similarly, the ‘summary statistics’ table also showed low expression of all detected genes in control4 **(Table 1)**. These QC results by iSeqQC indicated even though control4 sample was tightly clustered with its phenotypic group, due to its low-expression profile it could be considered as an outlier. A comparison of existing QC tools (sequencing and library) and iSeqQC is provided in **Table 2** indicating its importance in overall QC in expression-based sequencing experiments.

**Fig 2.**
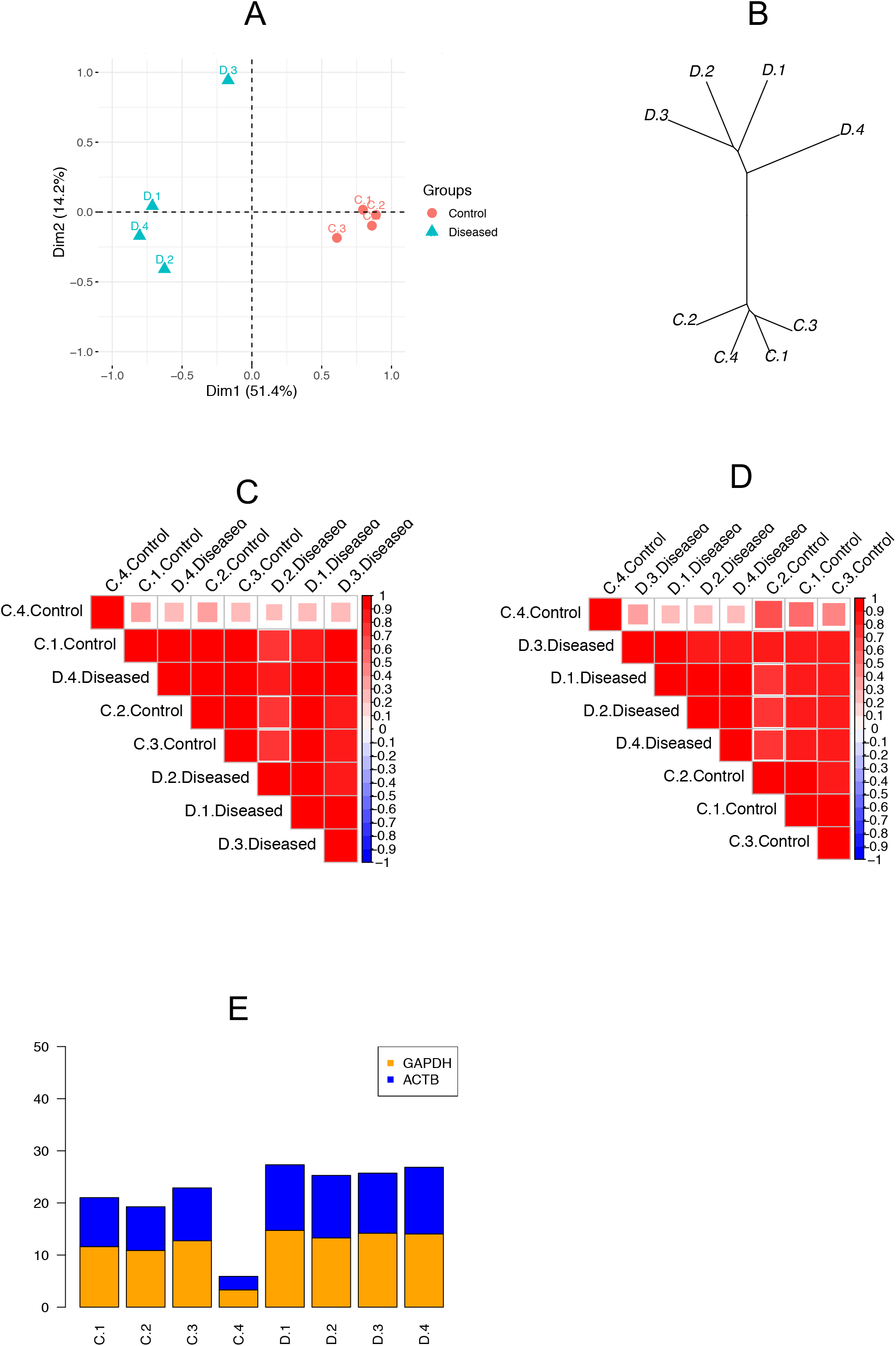
Quality control metrics produced by iSeqQC. A) Unsupervised PCA clustering showing tight cluster of samples within the phenotype; B) Hierarchical relationship assigning each sample to its own phenotypic cluster; C) Pearson correlation showing relationships between samples among biological replicates; D) Spearman correlation showing relationships between samples within their phenotype; E) Normalized expression of housekeeping genes (GAPDH and beta-actin) among different samples showing low expression of control4 sample.

**Table 1:** Summary statistics of control and diseased samples showing overall low expression of control 4 sample (iSeqQC output)

| **Sample**<br>**s**<br>**names** | **Numbe**<br>**r of**<br>**Genes** | **Mean** | **SD** | **Media**<br>**n** | **Mi**<br>**n** | **Max** | **Range** | **Skew** | **Kurtosi**<br>**s** |
| --- | --- | --- | --- | --- | --- | --- | --- | --- | --- |
| **D.1** | 24642 | 470.4<br>0 | 6087.7<br>8 | 74.13 | 0 | 633897 | 633897 | 71.49 | 38.78 |
| **D.2** | 24642 | 404.6<br>0 | 9244.1<br>1 | 57.82 | 0 | 122309<br>3 | 122309<br>3 | 103.4<br>4 | 58.89 |
| **D.3** | 24642 | 260.4<br>0 | 2769.2<br>0 | 48.93 | 0 | 289596 | 289596 | 63.11 | 17.64 |
| **D.4** | 24642 | 568.4<br>9 | 6646.3<br>3 | 100.82 | 0 | 685534 | 685534 | 69.23 | 42.34 |
| **C.1** | 24642 | 73.58 | 599.30 | 19.27 | 0 | 47932 | 47932 | 49.70 | 3.82 |
| **C.2** | 24642 | 61.91 | 459.74 | 22.24 | 0 | 46324 | 46324 | 72.30 | 2.93 |
| **C.3** | 24642 | 112.2<br>0 | 916.14 | 28.17 | 0 | 87803 | 87803 | 64.92 | 5.84 |
| **C.4** | 24642 | 12.92 | 16.79 | 8.90 | 0 | 829 | 829 | 11.25 | 0.11 |

**Table 2:** Features and capabilities of iSeqQC compared with other tools

| **Metrics** | **iSeqQC** | **QoRTs** | **QC3** | **RSeqQC** | **RNA-SeQC** |
| --- | --- | --- | --- | --- | --- |
| Summary expression | Yes | No | No | No | No |
| Dimensional reduction | Yes | No | No | No | No |
| Correlations | Yes | Yes | Yes | Yes | Yes |
| Housekeeping genes expression | Yes | No | No | No | No |
| Generate Plots | Yes | Yes | No | No | No |

Even though, iSeqQC flagged control4 to be an outlier, we decided to include it in further analysis for demonstration purposes. We performed differential expression analysis to obtain genes that are modulated in disease (absolute fold change≥2 and adjusted p value≤0.05) when compared to control. Here, we obtained 10,203 differentially expressed genes (DEGs), where 1278 were significantly up-regulated and 8925 were significantly down-regulated **(Fig. 3a)**. To access the impact of the outlier, we removed control4 sample (by changing sample phenotype file as shown in the workflow in **Fig. 4**) from the differential expression analysis and observed only 5311 DEGs, where 856 genes were significantly up-regulated and 4455 were significantly down-regulated **(Fig. 3b)**. This showed that if not removed, outliers can result in heavily biasing the data, which can result in false positive results.

**Fig 3.**
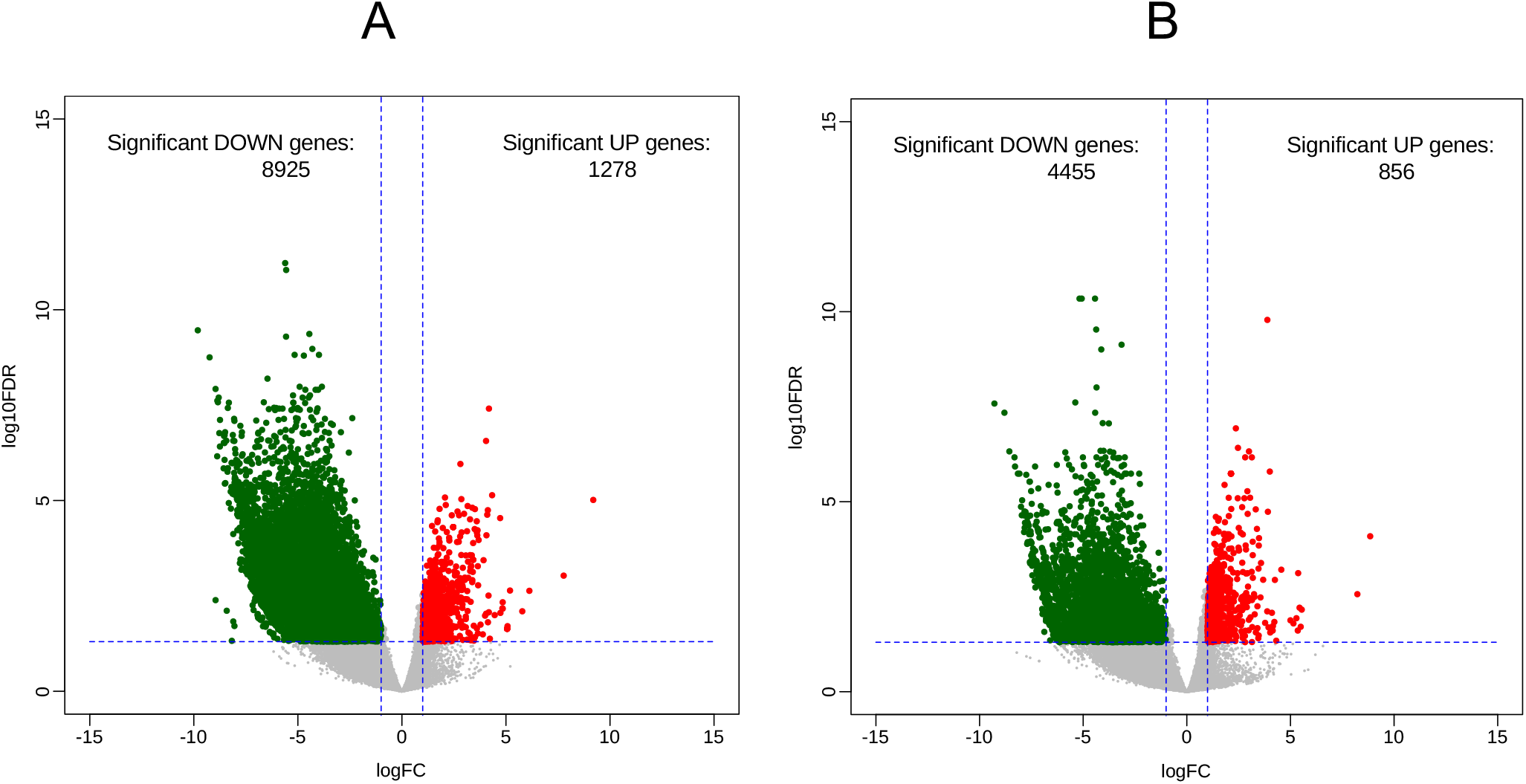
Change in number of differentially expressed genes before and after removal of outlier from differential expression analysis demonstrating the impact of an outlier in differential expression.

**Fig 4.**
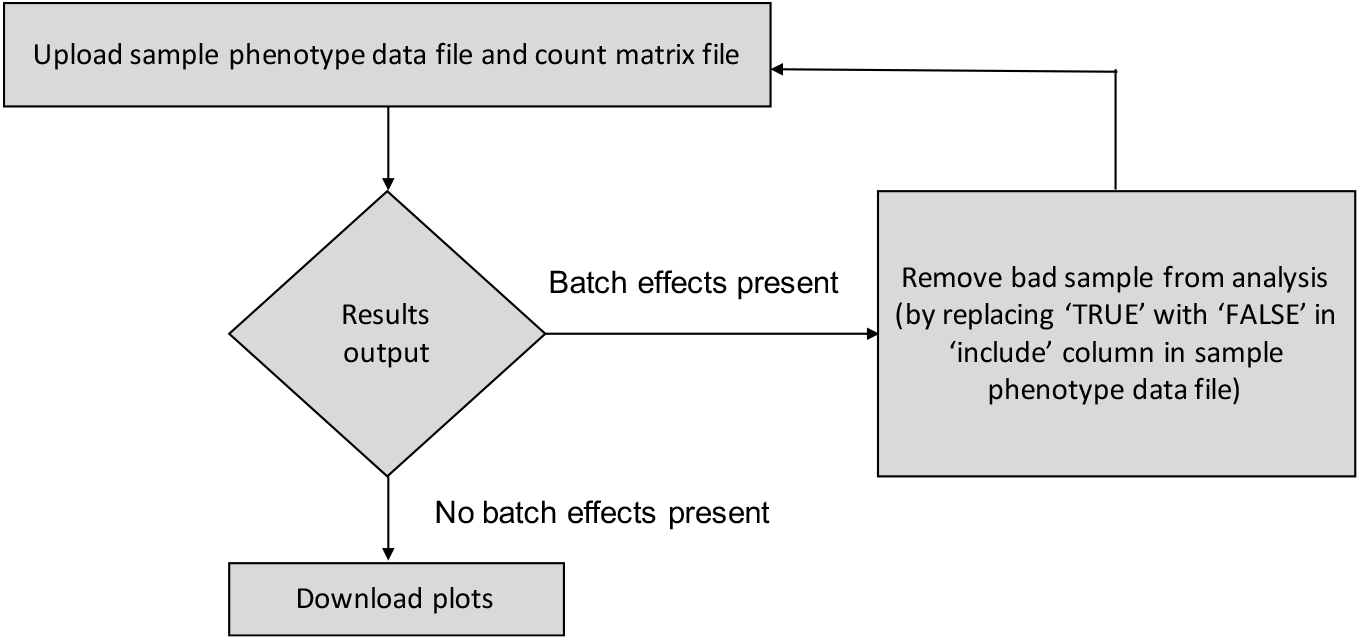
Workflow describing the steps to be followed to perform QC using iSeqQC

## DISCUSSION

Due to the complexity of high-throughput sequencing experiments, several phases of QC are required to identify any bias in the data. iSeqQC was designed to obtain comprehensive information on sample heterogeneity to detect outliers or cross-sample contamination in an expression-based sequencing experiment by implementing various statistical approaches including descriptive and dimensional reduction algorithms. iSeqQC was successful in identifying outliers in the studied dataset, which were missed by existing sequencing and library QC tools.

At present, there are no defined rules and tools available to perform QC on expression matrices for the detection of outliers in any sequencing experiment. The only approach is to use stand-alone statistical packages in R or other programming languages to compute PCA and correlations. However, as shown in the results these algorithms may not be sufficient to access the outliers. iSeqQC uses ensemble of various statistical methods to provide a detailed QC metrics in the form of a table and several graphical plots to identify any outliers. Additionally, at present while performing QC using stand-alone tools researchers have to re-run several lines of codes to re-evaluate the QC after removing the outliers. With iSeqQC, it is effortless as sample can be removed from the QC analysis by simply doing a minor change in sample phenotype sheet. Also, researchers spend hours generating publishable quality QC figures such as PCA and correlations plot, however iSeqQC by-default provides high-quality publication-ready figures.

While there exist several tools for assessing QC of sequencing experiments, each is limited to observe either sequencing and/or library quality. A few tools including QC3, QoRTs, RSeQC, and RNA-SeQC provide some information on outliers and cross-sample contamination but are not sufficient to provide in-depth sample qualities. QoRTs detects sample heterogeneity by analyzing read mapping, insert size distribution, cigar profile, and alignment clipping profile. RSeQC and RNA-SeQC uses Spearman and Pearson correlations to detect any outliers. QC3 is mainly focused to perform phase three QC only on Whole Exome Sequencing (WES) or Whole Genome Sequencing (WGS) data and does not include any quantitative sequencing technology such as RNA-seq. Additionally, all these tools require high-end computational resources and take hours to complete. As shown in the results these tools may not be sufficient to identify any outlier or cross-sample contamination in the complex dataset used here.

We acknowledge that due to the complexity of wet-lab protocols in sequencing technology, there are certain biasness that can evade the sample quality. Implementing statistical approaches gives only an idea of overall sample heterogeneity and is not sufficient to remove the samples from study. Use of additional methods such as Real time- Polymerase Chain Reaction (RT-PCR) is recommended to validate the findings.

## CONCLUSION

iSeqQC is a fast, light-weight, expression-based QC tool that detects outliers by implementing various statistical approaches. Implemented through web-interface and command-line interface, it generates high-quality publication-ready QC metrics for cross-comparison of samples.

## ACKNOWLEDGEMENTS

Authors thank Dr. Saul Surrey for his review and comments on the manuscript. This work was supported in part by an Institutional grant from the Sidney Kimmel Cancer Center of Thomas Jefferson University (NIH-NCI 2 P30 CA056036-23).

